# MR1 dependent MAIT cell activation is regulated by autophagy associated proteins

**DOI:** 10.1101/2022.09.20.508788

**Authors:** Prabhjeet Phalora, James Ussher, Svenja Hester, Carl-Philipp Hackstein, Emanuele Marchi, Jeffrey Y. W. Mak, David P. Fairlie, Paul Klenerman

## Abstract

The antigen presenting molecule, MR1, presents microbial metabolites to MAIT cells, a population of innate-like, anti-microbial T cells. It also presents an unidentified ligand to MR-1 restricted T cells in the setting of cancer. The cellular co-factors that regulate MR1 antigen presentation have yet to be fully defined. We performed a mass spectrometry-based proteomics screen to identify MR1 interacting proteins and discovered the selective autophagy receptor SQSTM1/p62. CRISPR-Cas9-mediated knock out of SQSTM1/p62 increased MAIT cell activation in the presence of *E.coli* but not the synthetic ligand 5-OP-RU whereas depletion of Atg5 and Atg7, key autophagy proteins, increased MAIT activation irrespective of the ligand used. This regulation appears to occur at an early step in the MR1 trafficking pathway. This data implicates distinct roles for autophagy associated proteins in the regulation of MR1 activity and highlights the contribution of the autophagy pathway to regulation of cellular antigen presentation.

## Introduction

Major Histocompatibility Related Protein 1 (MR1) is an evolutionary conserved, monomorphic, MHC like protein that presents microbial metabolites derived from vitamin B2 biosynthesis to a population of anti-bacterial innate-like CD8+ or double negative T cells termed Mucosal Associated Invariant T (MAIT) cells^1^. MAIT cells express high levels of CD161 and the semi-invariant TCR Vα7.2 and, upon activation, produce inflammatory cytokines such as IL17 and IFNγ as well as the cytolytic granules TNFα and Granzyme B^2–4^. The microbial ligands that bind to MR1 and activate MAIT cell effector functions are unstable pyrimidine intermediates, including 5-(2-oxoethylideneamino)-6-D-ribitylaminouracil (5-OE-RU) and pyrimidine-5-(2-oxopropylideneamino-6-D-ribitylaminouracil (5-OP-RU) formed by the condensation of the riboflavin biosynthetic precursor 5-amino-6-D-ribitylaminouracil (5-A-RU) with glyoxal and methylglyoxal, respectively^5–7^. In general, only microbes that generate riboflavin can activate MAIT cells^8^. However, the identification of MR1T cells, a novel population of T cells with diverse TCRs that are restricted by MR1 in the absence of bacteria, suggest that MR1 also presents non-microbial, endogenously derived ligands, including cancer specific antigens^9–12^. The recognition of, as yet, unidentified self-antigen(s) by MR1, as well as its non-polymorphic nature, make it an ideal candidate for the development of cancer immunotherapies and highlights the need for a better understanding of MR1 biology.

MR1 is expressed in most, but not all, mammalian species with a high degree of sequence conservation, particularly in the ligand binding domains^13^. Although MR1 mRNA is expressed in a variety of tissues and cell types, cellular protein levels and distribution appear to be tightly regulated and MR1 is difficult to detect at the cell surface, even after antigen exposure^14–16^. This contrasts with other MHC molecules which are constitutively expressed and readily detectable at the plasma membrane (PM). The mechanisms that govern this regulation remain poorly defined, although innate signalling molecules appear to be important^17,18^. Previous studies, mainly using over-expressed protein, have reported that the majority of MR1 is localized to the endoplasmic reticulum (ER)^19–22^, but it can also be found in the trans-Golgi network, endosomes and lysosomes^13,14,23^. ER resident MR1 is believed to be unbound and in an unfolded conformation. Upon interaction with its ligand, which forms a covalent bond with the sidechain of lysine 43, MR1 undergoes a conformational change that facilitates binding to β2M. This ternary complex exits the ER and traffics to the PM^19,24^. Little to no MR1 can be detected on the cell surface in the absence of antigen, which indicates that on the whole only ligand bound, fully folded MR1 can leave the ER. However, in some instances conformational changes leading to surface expression can occur in the absence of known ligands^17,25^.

Due to the inherently low expression of MR1, it has been difficult to resolve the cellular proteins, apart from β2M, that are involved in its trafficking from the ER to the PM to facilitate T cell activation^20–22^. Several groups have reported interactions between MR1 and components of the peptide loading complex (PLC), more commonly associated with MHC Class I trafficking^14^. This is supported by recent work from McWilliams et al, who demonstrated a key role for TPN and TAPBR in maintaining a pool of unbound MR1 in the ER, preventing its degradation before antigen exposure^26^. Additionally, loss of the ER associated P5-ATPase ATP13A1, a putative membrane transporter, reduced MR1 intracellular and surface protein levels^27^. After exiting the ER, certain Rab and SNARE proteins may be important for MR1 trafficking to the Golgi, as reported in the context of *M. tuberculosis* infection^23^. Once at the surface, MR1 is internalized and eventually degraded via a weak yet optimized interaction with the AP2 complex^28^.

Thus, the molecular machinery that governs MR1 cell surface expression and function has yet to be fully determined, and other unexplored pathways could be important. For instance, it was reported that MHC Class I internalization was regulated by autophagy^29^. Specifically, the removal of key autophagy proteins *in vivo* resulted in increased Class I expression at the PM, due to impaired receptor-mediated endocytosis and hence greater CD8+ T cell responses to viral infection. Autophagy is a key intracellular quality control pathway for the removal of damaged and defective cellular components. It is elevated in response to stress, which leads to the activation of mTOR kinase and formation of the autophagosome^30–32^. This double membrane structure elongates and matures to engulf the targeted material, which is then degraded upon fusion with a lysosome^33^. Each step of this process is carefully controlled by members of the Atg protein family^34^. Whether autophagy also contributes to regulation of MR1 cell surface expression and hence antigen presentation is unknown.

We aimed to identify the network of cellular proteins that interact with MR1 in bacterially exposed cells, by utilizing immunoprecipitation combined with mass spectrometry to perform a global, unbiased proteomics screen. Many of the proteins identified have known roles in antigen presentation, but several others, including components of the autophagy pathway, were also discovered. Thus, we sought to explore the role of autophagy and associated proteins in the regulation of MR1 expression and activity.

## Results

### A proteomics screen to define MR1 associated proteins during antigen presentation

Proteins that interact with MR1 may play an important role in its regulation as well as its function in antigen presentation. To identify these proteins, we used a Thp1 cell line stably expressing a HA tagged MR1 construct (Thp1.MR1.HA). Thp1 cells are a monocytic antigen-presenting cell (APC) line that express low levels of endogenous MR1 and can efficiently activate MAIT cells in a co-culture experiment. The tagged MR1 construct has been shown to be fully functional, akin to the wild-type protein^17^. Thp1 and Thp1.MR1.HA cells were incubated overnight with fixed *E. coli* and MR1 was immunoprecipitated using an antibody directed against the HA tag. Eluted samples were analyzed by liquid chromatography tandem mass spectrometry (LC-MS/MS) to identify cellular proteins that specifically associate with MR1 in the presence of antigen. As expected, β2M, a protein necessary for MR1 function, was identified as a significant MR1-interacting protein by this method, confirming the validity of this approach (Figure 1A and Supplementary Table S1). Several other proteins with known roles in antigen presentation and associated with the ER were also recovered, including Tap, Tapasin (TAPBP), Calnexin (CANX) and Calreticulin (CALR) (Figure 1A and 1B). In addition, proteins not previously reported to interact with MR1 were also identified, several of which have functions linked to ER degradation (SEL1L, AUP1, EDEM1) and autophagy (SQSTM1/p62, TMEM173/STING, TMEM59, LAMP-3) in particular. Interestingly, the alpha subunit of IL23 was identified as one of the proteins most highly associated with MR1. IL23 modulates the inflammatory response to infections via activation of the JAK-STAT pathway and contributes to several inflammatory and autoimmune diseases^35–37^. Overall, these data confirm earlier studies linking MR1 with proteins associated with antigen processing and presentation, particularly the PLC. However, they also demonstrate that the cellular network of MR1 interacting proteins may be more expansive than previously described.

**Figure 1:**
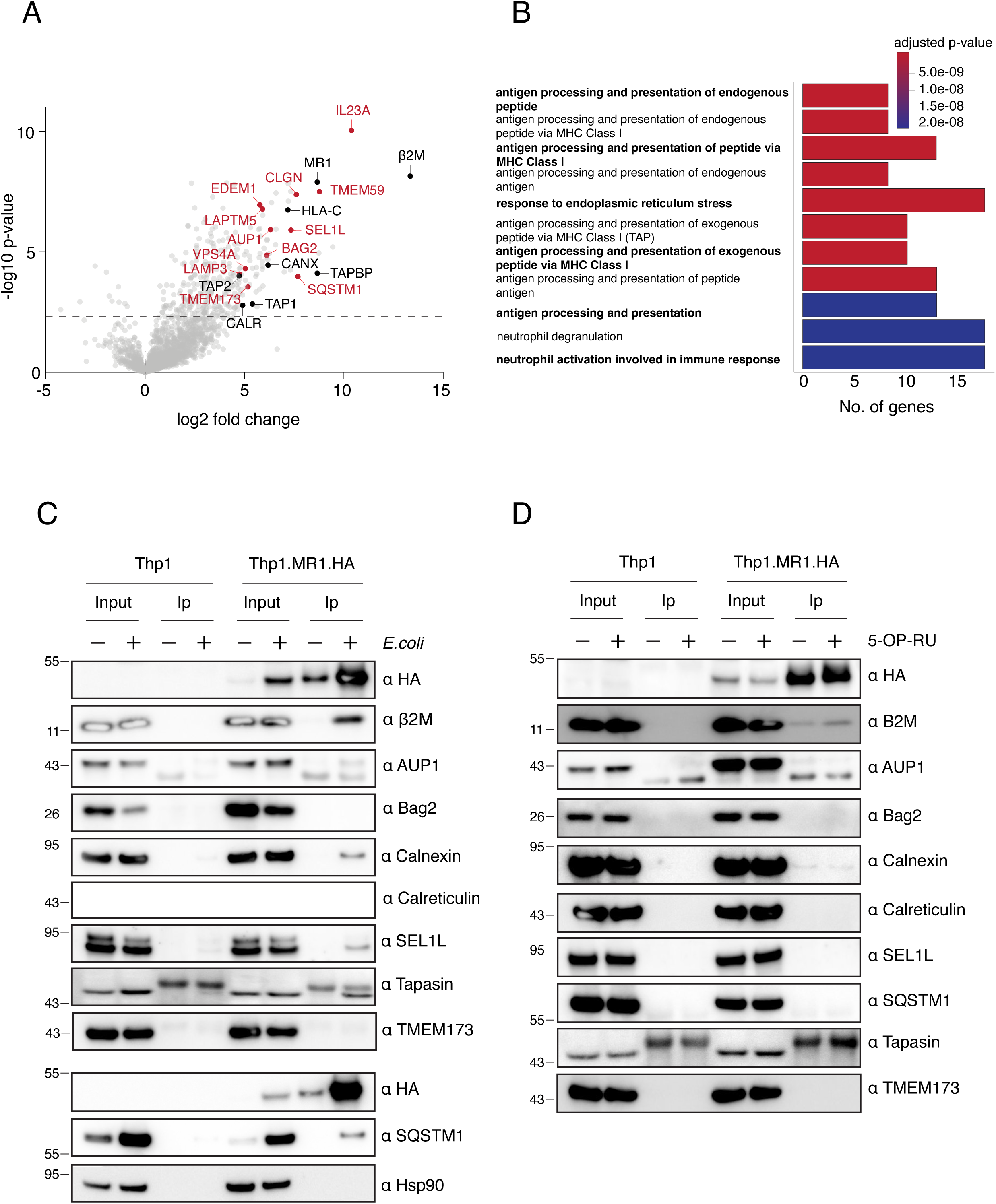
A proteomics screen to define MR1 associated proteins during antigen presentation. A. Thp1 and Thp1.MR1.HA cells were incubated with fixed *E. coli* overnight and lysed in 0.5% NP40. MR1 was immunoprecipitated from lysates using a HA antibody conjugated to magnetic beads. Bound proteins were eluted, trypsin digested and analyzed using LC-MS/MS. Significantly enriched proteins in the presence of MR1 are shown in the top right quadrant. Components of the peptide loading complex are highlighted in black and other proteins of interest are highlighted in red. Data were combined from 2 independent experiments each consisting of 3 technical repeats and were analyzed using MAXQuant software. B. Pathways analysis of the most highly enriched genes in the MR1 fraction. C. Thp1 and Thp1-MR1.HA cells were incubated with fixed *E. coli* overnight and then lysed in 0.5% NP40. MR1 was immunoprecipitated using a HA antibody conjugated to magnetic beads. Immunoprecipitates were analyzed by western blot and probed with the indicated antibodies. D. As for C but cells were incubated with 5-OP-RU overnight. Data in C and D are representative of at least 3 independent experiments. The position of molecular weight marker bands (kDa) are indicated to the left of the immunoblots.

### Ligand dependent cytokine induction may alter the composition of the MR1 proteome

Next, we sought to validate the interactions between MR1 and several of the proteins identified by the proteomics screen. Immunoprecipitated MR1.HA, from Thp1.MR1.HA cells incubated with fixed *E. coli* overnight was run on SDS-PAGE gels and then probed with specific antibodies targeting the proteins of interest. β2M was efficiently co-immunoprecipitated with MR1 by this method. Components of the PLC, including Calnexin, Tapasin and Calreticulin were found to associate with MR1 (Figure 1C). SEL1L and SQSTM1, which were identified by the mass spectrometry analysis, were also pulled down in the presence of MR1 but IL23A and TMEM173/STING were not (Figure 1C and Supplementary Figure 1). To demonstrate the specificity of the interactions, a control protein not identified by the screen, HSP90, also failed to associate with MR1. When the co-immunoprecipitations were performed in the presence of the synthetic MR1-binding ligand 5-OP-RU, rather than *E. coli*, these interactions could no longer be recapitulated, except for β2M and, more weakly, Calnexin, both in the presence and absence of antigen (Figure 1D). Reducing the incubation period from overnight to 2 hours still did not recover all of the interactions as observed with *E. coli* (Supplementary Figure 2). These results suggest that other components of *E. coli* may be affecting engagement with cellular proteins as has been previously reported^11,16,38^. This may be supported by the observation that co-incubation of Thp1.MR1.HA cells with fixed *E. coli* leads to an increase in MR1.HA (and to a lesser extent SQSTM1) protein expression, even at lower bacteria to cell ratios (Supplementary Figure 3A), possibly due to changes in the cellular cytokine environment that promotes MR1 expression and/or stabilization. This was not observed when the cells were treated with synthetic 5-OP-RU alone (Supplementary Figure 3B), even in the presence of the cytokines IL12 and IL18. Incubation with 5-OP-RU and IL12/18 did not result in a detectable interaction between MR1 and SQSTM1 (Supplementary Figure 3C). Further, use of IL12 and IL18 blocking antibodies did not prevent an increase in MR1 expression when the cells were also incubated with *E. coli* (Supplementary Figure 3D), indicating that other cytokines/bacterial interactions must also be important. Thus, the composition of the MR1 proteome may differ according to the nature of the ligand present and may also be modulated by cytokine responses.

### Knock-out of the autophagy adaptor protein SQSTM1 leads to increased MAIT cell activation in the presence of E. coli

We focused on the autophagy associated protein SQSTM1 (also known as p62) because in the presence of *E. coli* its expression levels increased and it consistently interacted with MR1 in our system. SQSTM1 is an adaptor molecule that binds to ubiquitinated cargo in the ER through its UBA domain and LC3 through its LIR domain, to target proteins to autophagosomes for degradation via selective autophagy^39,40^. It also has roles in several cellular signaling pathways^41,42^. To understand the functional significance of the MR1-SQSTM1 interaction, we utilized CRISPR-Cas9 technology to knock out SQSTM1 (and β2M as a positive control) from wild-type Thp1 cells (expressing endogenous levels of untagged MR1) and established single-cell clones by limiting dilution. All clones were verified by immunoblot analysis and sequencing (Figure 2A). These cells were then incubated overnight with either DMSO, fixed *E. coli,* 5-OP-RU or the non-MAIT stimulatory synthetic ligand Ac-6-FP and levels of surface MR1 were compared. Both β2M knockout clones (B1 and B6), showed substantially reduced MR1 surface staining in all conditions tested, as expected (Figures 2B and 2C). However, the results from the SQSTM1 knockout clones (S1, S2, S3 and S4) were more variable (Figures 2B and 2C). There were no significant differences between the clones and the control cell line in the *E. coli* condition, however two of the clones (S1 and S4) displayed significantly reduced MR1 surface levels in the presence of 5-OP-RU and Ac-6-FP. Surface MHC Class I levels were mostly not affected in these clones (Supplementary Figure 4).

**Figure 2:**
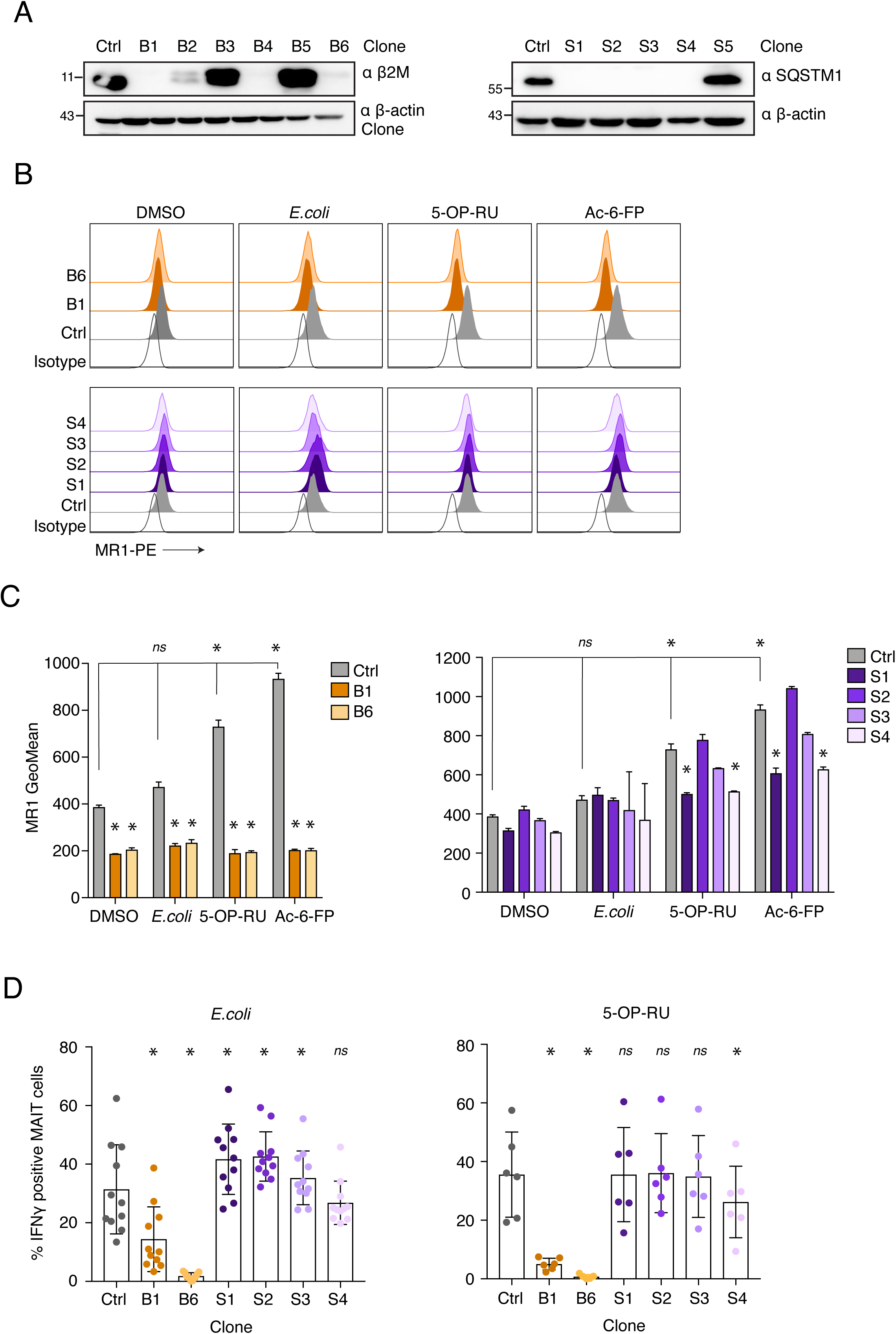
Knock-out of the autophagy adaptor protein SQSTM1 leads to increased MAIT cell activation in the presence of *E. coli*. A. Thp1 cells were transduced with gRNAs directed against β2M (left) and SQSTM1 (right). Single cell clones were established by limiting dilution and knockout confirmed by western blot by probing with the indicated antibodies. The position of molecular weight marker bands (kDa) are indicated to the left of the immunoblots. B. Knockout clones for β2M (B1 and B6) and SQSTM1 (S1, S2, S3 and S4) produced in A were incubated with DMSO, fixed *E. coli*, 5-OP-RU or Ac-6-FP overnight and then stained for surface MR1 expression. Representative histograms displaying geometric MFIs are shown. C. Quantification of data as described in B. Displayed is the average from 3 independent experiments ± SD. D. Knockout clones for β2M and SQSTM1 as described in A were incubated with either fixed *E. coli* (left) or 5-OP-RU (right) overnight. Cells were incubated with CD8+ enriched T cells from healthy donors for 5 hours with Brefeldin A added for the final 4 hours. % IFNγ production ± SD from CD161++Vα7.2+ cells was determined by FACS. For C, data were acquired from 11 donors across 3 experiments, for D, data were acquired from 6 donors across 2 experiments. For C and D, statistical significance was calculated using a one-way ANOVA, * = p < 0.05, ns = p > 0.05.

To assess whether these changes impacted MR1 antigen presentation, we performed an assay to measure levels of MAIT cell activation. β2M and SQSTM1 knock out clones were incubated with *E. coli* or 5-OP-RU overnight and then co-cultured with purified CD8+ T cells for 5 hours. IFNγ production from the CD161++ Vα7.2+ population was measured by FACS (Figure 2D). In cells depleted of β2M there was a significant reduction in MAIT cell activation in both the *E. coli* and 5-OP-RU conditions (Figure 2D). In contrast, we observed a significant increase in IFNγ production from MAIT cells after incubation with the SQSTM1 knock out cells in the presence of *E. coli.* Interestingly, in the presence of 5-OP-RU, MAIT activation was either unaffected or reduced compared to the control (Figure 2D, right panel). A second set of clonal cells derived from a different guide RNA targeting SQSTM1 also showed an increase in MAIT cell activation following *E. coli* stimulation and a significant decrease with 5-OP-RU stimulation (Supplementary Figures 5A and 5B). The only exception to this was clone #5, which showed a significant decrease in the presence of *E. coli* (Supplementary Figure 5B). This clone grew substantially slower than the others, which may explain its contrasting phenotype. The increase in MAIT cell activation in the presence of *E. coli* was also observed in the bulk population of cells before generation of the clonal lines and was blocked with the use of an anti-MR1 antibody (Supplementary Figures 6A and 6B). These results could also be reproduced when SQSTM1-depleted C1R cells, a B cell line, were used as the APC instead of Thp1 cells (Supplementary Figures7A and 7B). Overall, in the absence of SQSTM1, MAIT cell activation was increased but only in the presence of *E. coli* and not 5-OP-RU. This implies that additional cellular changes induced by *E. coli* affect MR1 expression, stabilization and/or function.

### Depletion of Atg5 and Atg7 increases MAIT cell activation

The varied results observed with knockout of SQSTM1 on MAIT cell activation in the presence of fixed *E. coli* versus 5-OP-RU could be attributable to the multiple other cellular functions of this protein, distinct from autophagy^41,43–45^. Additionally, although SQSTM1 is an important component of selective autophagy, its role may be functionally redundant with other similar adaptor proteins^40,46^. To determine the effects of autophagy more precisely on MRI function we targeted two essential components of the autophagy machinery. Atg5 and Atg7 have significant roles in the formation of the autophagosome and are routinely depleted *in vitro* and *in vivo* to study the effects of autophagy. Reduced expression of these proteins in wild-type Thp1 cells by CRISPR –Cas9 treatment (Figure 3A) led to impaired autophagy in these cells as assessed by the reduction in autophagic flux in both normal and amino acid starved cells^47^ (Supplementary Figure 8). Surface MR1 levels in Atg5 and Atg7 CRISPR-Cas9 treated cells were moderately increased after incubation with *E. coli* and 5-OP-RU compared to control cells, although these differences were not statistically significant (Figure 3B). In addition, IFNγ production by MAIT cells was also increased with both ligands after co-culture with these cell lines (Figures 3C and 3D). This was also true when C1R cells were used as the APC rather than Thp1s (Supplementary Figure 9). The use of multiple guide RNAs against two separate proteins in the autophagy pathway with similar outcomes means that it is less likely that the observed phenotypes are due to off-target effects.

**Figure 3:**
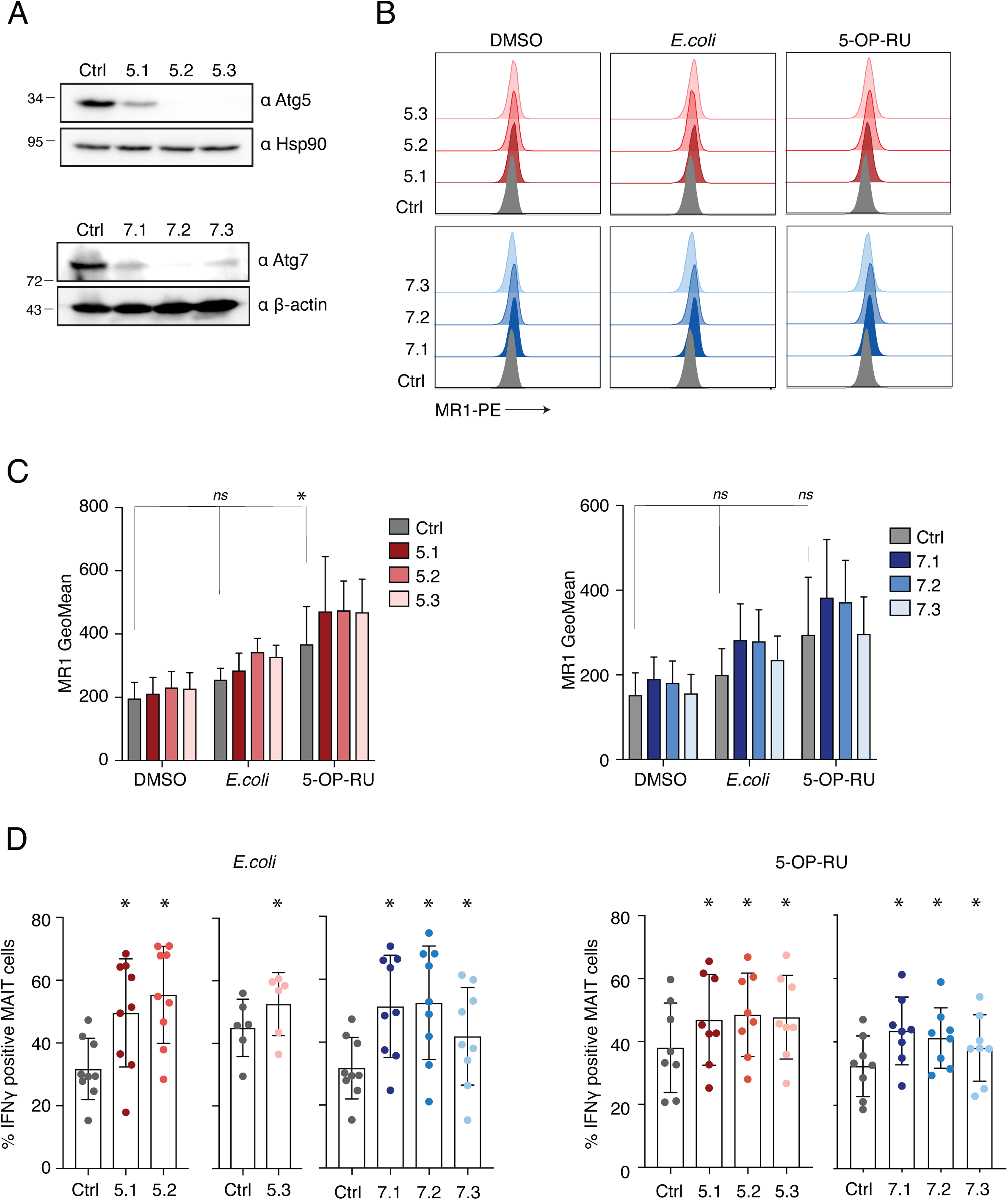
Depletion of Atg5 and Atg7 increases MAIT cell activation. A. Thp1 cells were transduced with gRNAs targeting Atg5 (left) or Atg7 (right). Protein depletion was analyzed by western blot. The position of molecular weight marker bands (kDa) are indicated to the left of the immunoblots. B. Atg5 (top) and Atg7 (bottom) depleted cell lines as produced in A were incubated with DMSO, fixed *E. coli* or 5-OP-RU overnight and then stained for MR1 surface expression. A representative histogram is shown on the left, with geometric MFIs displayed, and the average ± SD of at least 3 independent experiments is shown in the bar plots on the right. Cells described in A were incubated with *E. coli* (C) or 5-OP-RU (D) overnight and analyzed for MAIT cell IFNγ production as described in Figure 2D. The data for Atg 5.3 is displayed separately as the experiments for this cell line were performed at a later timepoint using different donor cells. For C, data were acquired from 9 donors across 3 experiments (except for Atg5.3, where data was acquired from 6 donors across 2 experiments) and for D, data were acquired from 8 donors across 2 experiments. For B, C and D statistical significance was calculated using a one-way ANOVA, * = p < 0.05.

As surface MR1 levels were not significantly increased in the autophagy-deficient cells, we generated Thp1.MR1.HA cells transduced with Atg5 and Atg7 gRNAs to check total protein levels by western blot. MR1 protein expression was higher in the Atg-depleted cells compared to the control cells, even at baseline (Supplementary Figure 10). The same was true for SQSTM1, a protein known to be regulated by autophagy, whereas other ER resident proteins remained unchanged. We also sought to determine if any of the interactions we previously found with MR1 (Figure 1C) remained intact in the absence of autophagy. Detectable interactions between MR1 and β2M, SQSTM1, Calnexin and Tapasin were still observed in the knockdown cell lines (Supplementary Figure 10). Taken together, these results demonstrate that MR1 protein levels are upregulated in autophagy-deficient cells intracellularly and to a lesser extent at the cell surface. These changes in protein expression have a significant impact on MR1 antigen presentation and subsequently MAIT cell activation, implicating a role for the autophagy pathway in the regulation of MR1 activity.

### Manipulation of autophagy alters MR1 intracellular and surface expression levels

As a complementary approach we also used a panel of chemical reagents to manipulate the autophagy pathway. 3MA and wortmannin are PI3K inhibitors which prevent activation of mTOR and thus block autophagosome formation at an early stage^30,47^. Treatment of Thp1.MR1.HA cells with these inhibitors led to an increase in total MR1 protein levels as determined by western blot (Figure 4A). Amino acid starvation of cells induces autophagy and subsequent treatment with bafilomycin prevents the autophagosomes from fusing with lysosomes and being degraded. Incubation of Thp1.MR1.HA cells in EBSS media, which lacks amino acids, slightly increased MR1 levels but was not further enhanced with subsequent addition of bafilomycin (Figure 4A). To further assess the effects of these reagents on MR1 expression and localization we imaged HeLa cells stably expressing MR1.HA. HeLa cells were used in this case as they are more amenable to imaging methods than Thp1 cells and are routinely used to study MR1 for this purpose^19^. HeLa.MR1.HA cells were able to efficiently activate MAIT cells in a co-culture experiment and this could be blocked with the use of an anti-MR1 antibody (Supplementary Figure 11). Treatment of HeLa.MR1.HA cells with both autophagy inhibitiors (3MA, wortmannin and bafilomycin) and inducers (EBSS) led to a significant increase in the average number of SQSTM1 foci per cell (Figure 4B), as previously reported^48,49^. Additionally, treatment with 3MA led to a significant increase in MR1 cytoplasmic signal intensity compared to the DMSO control (Figure 4B). Following on from this we specifically looked at MR1 surface levels in human primary monocytes and Thp1.MR1.HA cells. Surface MR1 levels were significantly increased in primary monocytes with the addition of bafilomycin and wortmanin and significantly decreased following EBSS treatment (Figure 4C). Similar trends were also observed in Thp1.MR1.HA cells. Pre-incubation with 5-OP-RU before addition of the autophagy inhibitors significantly elevated MR1 surface expression compared to the control condition in both cell types. Surprisingly, this increase was also seen when autophagy was enhanced with the use of EBSS (Figure 4C, right graphs), indicating that the addition of antigen can rescue MR1 bound for degradation via autophagy. Therefore, these data demonstrate that perturbations in cellular autophagy can affect both intracellular and surface levels of MR1 expression.

**Figure 4:**
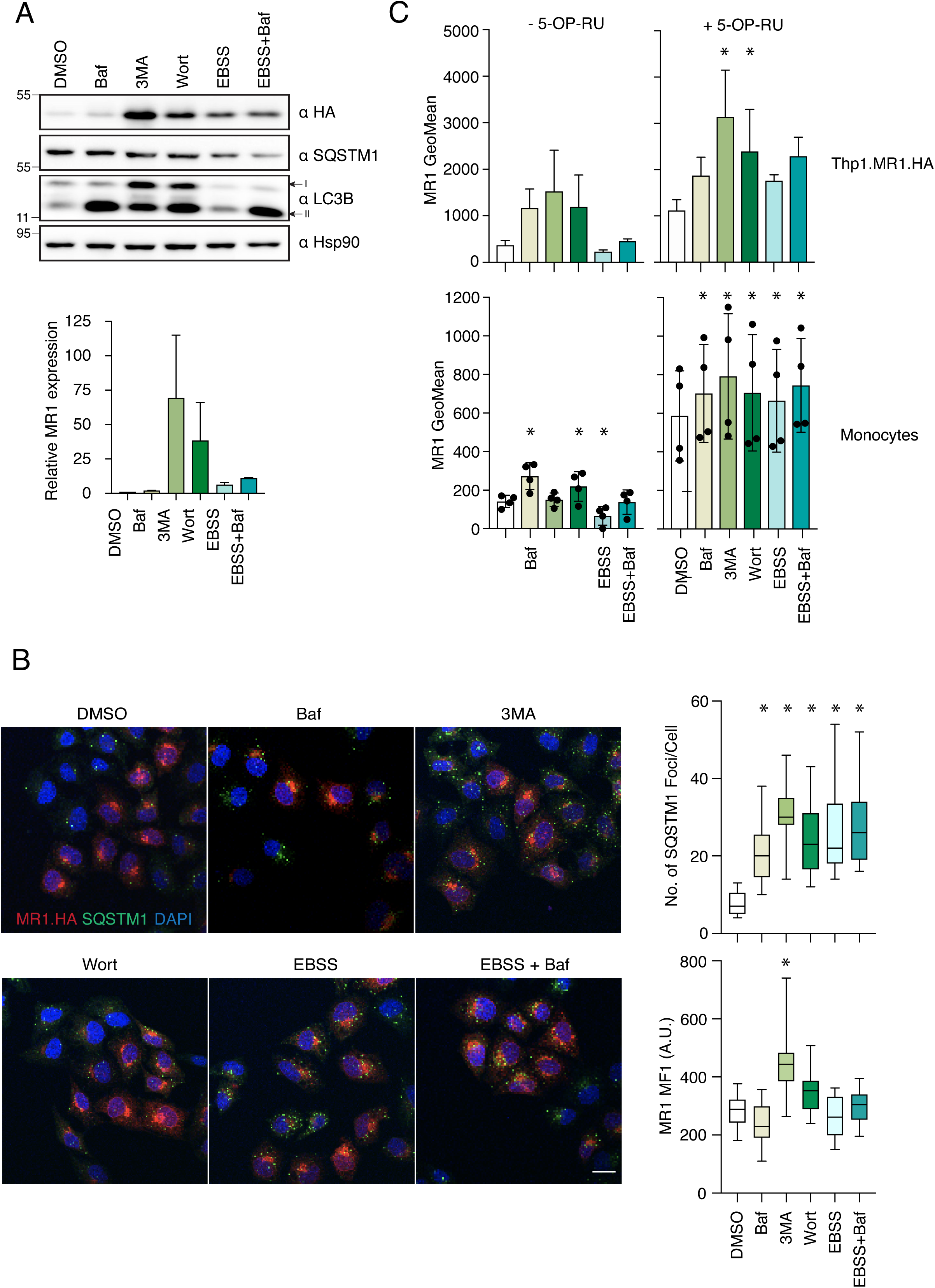
**Manipulation of autophagy alters MR1 intracellular and surface expression levels**. A. Thp1.MR1.HA cells were incubated with DMSO, 100nM bafilomycin (Baf), 5mM 3MA, 1μM wortmannin (Wort), EBSS or EBSS + 100nM bafilomycin (EBSS+Baf) for 4 hours (bafilomycin was added for the final 2 hours) and then lysed. Cell lysates were probed with the indicated antibodies by western blot. The position of molecular weight marker bands (kDa) is indicated to the left of the immunoblots and the LC3-I (I) and LC3-II (II) bands are indicated on the right. Image is representative of 2 independent experiments with the average ± SD displayed in the bar graph. B. HeLa cells expressing MR1.HA were grown on coverslips and incubated in normal media +/- Baf, 3MA or Wort and EBSS media +/- Baf for 4 hours with bafilomycin added for the final 2 hours. Cells were fixed and then permeabilized with 0.1% TritonX-100 and stained with anti-HA and anti-SQSTM1 primary antibodies and species-specific Alexa Fluor conjugated secondary antibodies. 40 cells were quantified across 2 independent experiments for the number of SQSTM1 foci per cell (top graph) and Mean Fluorescence Intensity (MFI) of MR1 (bottom graph). Data are represented in a box and whisker plot displaying Q1, median and Q3 as well as the minimum and maximum values. Scale bar = 10μm. C. Thp1.MR1.HA cells (top) or primary human monocytes (bottom) were pre-incubated (right) or not (left) with 5-OP-RU for 2 hours and then treated with autophagy reagents as in A and stained for surface MR1 expression. Displayed is the average ± SD from 3 independent experiments (top) or from 4 healthy blood donors across 2 experiments (bottom). Statistical significance was calculated using a two-way ANOVA, * = p < 0.05.

**Figure 5:**
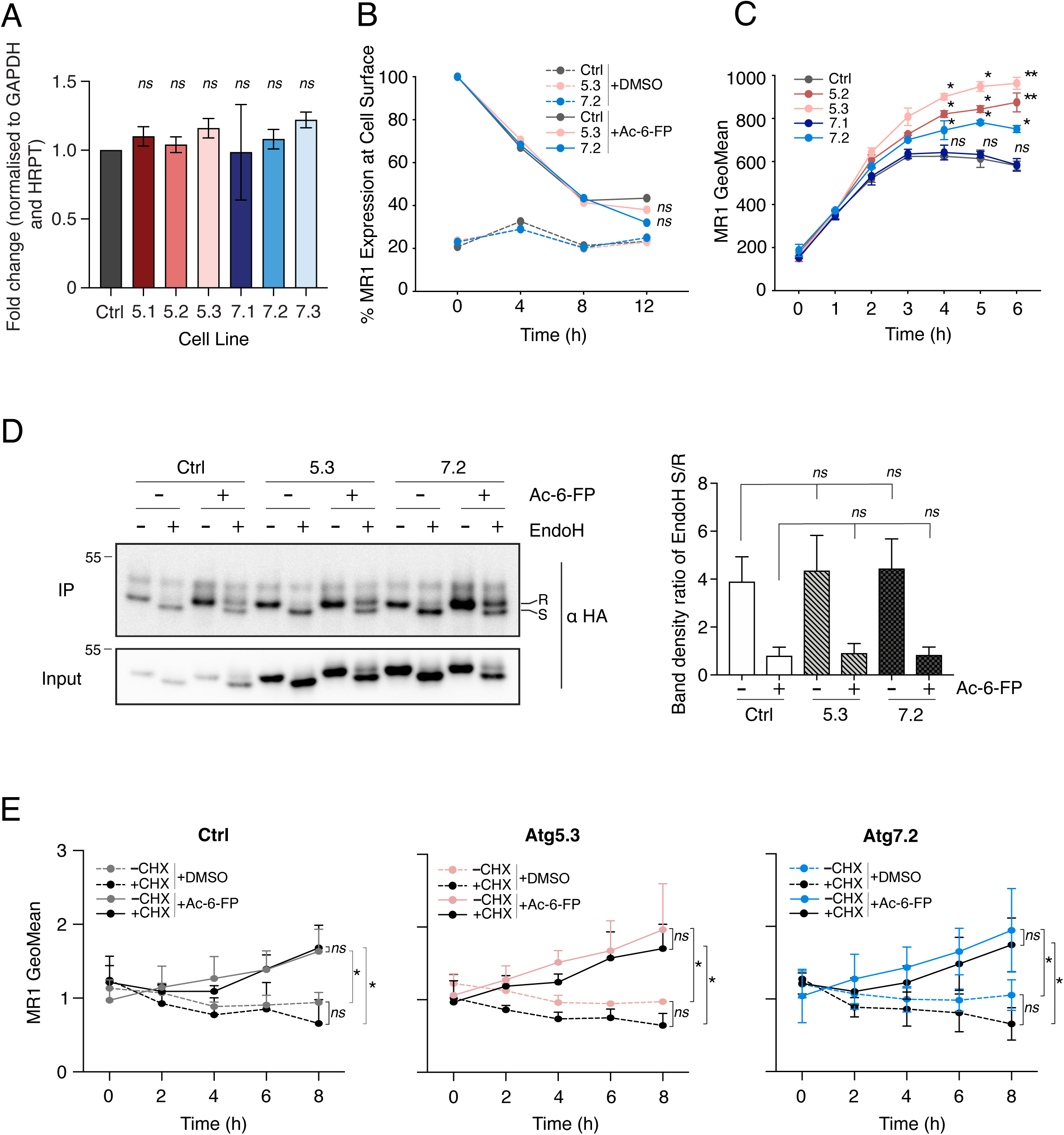
Autophagy regulates MR1 at an early step of the MR1 trafficking pathway. A. MR1 transcript levels were measured in Atg5 and Atg7 depleted cells and fold change in expression compared to the control cell line was calculated. ΔΔCT values were normalized to the housekeeping genes, GAPDH and HRPT. Data is the average of 2 independent experiments ± SD, each consisting of 3 technical repeats. B. Control or Atg5 (5.3) and Atg7 (7.2) depleted cell lines were incubated overnight with either DMSO or Ac-6-FP. Cells were washed and incubated in ligand free media for up to 12 hours. Surface MR1 levels were measured at the indicated timepoints and are displayed as % expression compared to the 0 hour timepoint + Ac-6-FP which is set at 100% for each cell line. Data is the average of 2 independent experiments. C. Control or indicated Atg5 (5.2 and 5.3) and Atg7 (7.1 and 7.2) depleted cell lines were incubated with Ac-6-FP and surface MR1 levels were measured every hour for up to 6 hours. Displayed are the average gMFIs ± SD from 3 independent experiments. D. Thp1.MR1.HA control or Atg5 (5.3) or Atg7 (7.2) depleted cell lines were incubated with DMSO or Ac-6-FP overnight. Cells were lysed and MR1 immunoprecipitated using a HA antibody conjugated to magnetic beads. Samples were treated or not with EndoH for 1.5 hours and analyzed by western blot by probing with the indicated antibodies. Input samples are shown as a control below. The position of the molecular weight marker bands (kDa) are indicated to the left of the immunoblots and the MR1 EndoH resistant (R) and senstitive (S) bands are indicated on the right. Image is representative of 4 independent experiments with the quantification from all immunoblots shown in the bar plot below. Data is presented as the average ratio of S/R ± SD for the EndoH treated samples only. E. Control or Atg5 (5.3) and Atg7 (7.2) depleted cell lines were incubated with 20μM cycloheximide (CHX) for 8 hours, with DMSO (dotted lines) or Ac-6-FP (solid lines) added after the 1st hour. Cells were stained for MR1 surface expression every 2 hours and displayed is the average gMFI ± SD from 3 independent experiments. Statistical significance for all experiments was analyzed using a two-way ANOVA, * = p<0.05, ns = p > 0.05.

### Autophagy regulates pre-existing pools of MR1, possibly before it reaches the plasma membrane

Although the data presented thus far implicated the autophagy pathway in the regulation of MR1 function, it was unclear at which step of MR1 trafficking, from the ER to the plasma membrane, this regulation was occurring. MR1 transcripts were comparable between control and autophagy-depleted cells (Figure 5A), suggesting that this effect was not due to differences in the levels of mRNA. MR1 at the cell surface may be internalized and either degraded or recycled via an autophagic pathway as described for MHC Class 1. To address this point, we incubated control and autophagy-depleted cells with DMSO or Ac-6-FP overnight and then chased in ligand free media for up to 12 hours (Figure 5B). As the knock down cell lines behaved similarly in terms of MR1 phenotype we used Atg5.3 and Atg7.2 as representative examples for this and subsequent experiments. For each cell line the 0h timepoint (beginning of the chase) plus Ac-6-FP was set to 100% to facilitate direct comparisons. We found no difference between the cell lines in the rate of decline of surface MR1 over the 12-hour chase period. Thus, MR1 already at the cell surface is not differentially regulated by autophagy. In contrast, differences in surface MR1 expression in control compared to Atg5 and Atg7-depleted cells were observed as early as 2 hours post ligand addition and become more pronounced at the 6-hour time point (Figure 5C). This indicates that regulation by autophagy occurs early in the trafficking of MR1, potentially before it reaches the plasma membrane. As MR1 is known to traffic via the Golgi to reach the plasma membrane, we sought to determine the amounts of EndoH sensitive (pre-Golgi processed) to EndoH resistant (Golgi processed) MR1 forms in control and autophagy-deficient cells (Figure 5D). There appeared to be no difference in the ratio of EndoH sensitivity between the cell lines, demonstrating that MR1 is processed normally through the Golgi apparatus in the absence of autophagy. Lastly, we treated control and Atg-deficient cells with the protein synthesis inhibitor Cycloheximide (CHX) in the presence and absence of synthetic ligand to determine whether autophagy is regulating new or pre-existing MR1 protein populations (Figure 5E). The level of upregulation of surface MR1 in Atg depleted cells, as well as control cells, was the same in CHX treated and untreated conditions suggesting that MR1 surface expression is not dependent on new protein synthesis. Thus, Atg regulation is potentially occurring on pre-existing pools of MR1, the majority of which, in the absence of antigen, is contained within the ER^19^.

## Discussion

MR1 plays an essential role in presenting microbially derived antigens to MAIT cells and, some additional recently-defined antigens to MR1T and cancer-targeting T cells^11,50,51^. These functions are important in anti-microbial immunity and in the specific recognition of transformed cells, respectively, as demonstrated by the high degree of sequence conservation of this protein^52^. Although the functions of MR1 are becoming clearer, how these functions are controlled by cellular co-factors remains to be fully characterized.

In this study, we were able to identify and validate several existing and newly identified MR1 interacting proteins. Some of these proteins have known roles in antigen presentation, particularly components of the PLC. This complex is more commonly associated with MHC Class I presentation and thus supports earlier and more recent findings that MR1 trafficking utilizes a similar pathway^1,21,26^.

We used a HA tagged overexpressed MR1 construct to obtain sufficient protein to perform the mass spectrometry analysis. We previously demonstrated that the tag did not interfere with the correct folding and function of MR1^17^. Thp1 cells are natural APCs and thus the cellular components required for MR1 antigen presentation are fully active and present at endogenous levels. Further, we used fixed bacteria as well as synthetic 5-OP-RU and Ac-6-FP, as the ability of the chemically synthesized antigens to access the MR1 compartment may not be fully recapitulated under *in vivo* conditions. This experimental setup was validated by recovery of β2M as a major MR1 interacting protein. We also verified the interaction of MR1 with previously unidentified proteins such as SEL1L and SQSTM1. However, other interactions could not be confirmed by co-immunoprecipitation, most notably with IL23A. Although this protein could be easily detected in cells, there was no association with MR1 in our validation experiments. This interaction may not be amenable to analysis by co-immunoprecipitation and other methods may need to be employed to determine its validity.

Although we identified interactions between MR1 and components of the PLC, including TAP2, this was only possible in the presence of *E. coli* and not at baseline or with 5-OP-RU alone, even with reduced incubation times to account for 5-OP-RU instability^7^. This contrasts with the work of McWilliams et al, who performed a similar proteomics screen and were able to recover several of these interactions in the absence of ligand in C1R and Thp1 cells and in the presence of Ac-6-FP in C1R cells^26^. The reason for this discrepancy remains unclear but may reflect differences in experimental conditions, for example the type of tagged MR1 construct used, the level of overexpression, and the use of Ac-6-FP versus 5-OP-RU.

Despite these differences, both proteomics screens, and another recently conducted in BEAS2B cells^53^, identified SQSTM1 as an MR1 interacting protein. SQSTM1 plays an essential role in the selective autophagy pathway as it facilitates the removal of ubiquitinated proteins as well as bacteria from cells via autophagosomes^39,49,54–56^. In the presence of *E. coli,* knockout of SQSTM1 did not affect MR1 surface expression but did increase antigen presentation to MAIT cells in nearly all clones tested. However, in the presence of 5-OP-RU loss of SQSTM1 either reduced MAIT activation or had no effect, implying that in certain circumstances, SQSTM1 may be required for efficient MR1 antigen presentation and that regulation of MR1 may be antigen specific, as has previously been suggested^16,17,23,38^. Further studies are required to determine whether this specificity has a biological significance, for example, in response to intracellular versus extracellular bacteria.

In contrast to these results, depletion of the autophagy specific molecules, Atg5 and Atg7, led to increased MAIT activation in the presence of both *E. coli* and 5-OP-RU. Total MR1 protein levels were also increased when cells were treated with the autophagy inhibitors 3MA and wortmannin. Use of the late-stage autophagy inhibitor bafilomycin resulted in an increase in surface MR1 levels in primary monocytes and Thp1 cells, although intracellular protein expression was not affected. Thus, when the generation of autolysosomes is blocked, surface MR1 accumulates, implicating autophagy mediated degradation in the regulation of MR1 protein levels.

Although the effect sizes observed here are small, the physiological impact of increased MAIT activation may be more significant and requires further exploration. The highly regulated nature of MR1 expression, even after antigen exposure, strongly suggests that activation of MAIT cells is finely balanced and any perturbations could have important repercussions for innate immune responses.

The role of autophagy in antigen presentation has long been recognized. MHC-II presentation to CD4+ cells is dependent upon peptides generated from degraded cellular constituents and processed via the autophagy pathway. Autophagy may also contribute to Tap independent presentation of viral epitopes for MHC-I presentation. Aside from antigen processing, autophagy associated proteins have been shown to control levels of MHC Class I and CD1 on the surface of dendritic cells through the adaptor proteins AAK^29^ and AP2^57^ respectively. Interestingly, AP2 was recently identified as the receptor controlling optimal rates of MR1 internalisation^28^. Thus, one possibility is that inhibition of autophagy prevents the degradation of surface MR1, maintaining more protein at the plasma membrane and hence greater MAIT activation. Although we found no difference in the rate of removal of MR1 from the cell surface in control versus knock down cells following antigen exposure, closer analysis of this step of the MR1 lifecycle will be required to determine whether AP2 mediated autophagic degradation is involved in MR1 regulation.

An alternative possibility is that MR1 is degraded by autophagy at an earlier step in the trafficking pathway. In support of this, induction of autophagy through amino acid starvation led to a decrease in surface MR1 levels, which were dramatically elevated in the presence of 5-OP-RU. This shows that MR1 targeted for degradation can be rescued by the addition of antigen which implies that it is the population of unbound, but ligand receptive, MR1 that is regulated by autophagy. This regulation may occur through a process such as ER associated degradation (ERAD), a quality control system which removes misfolded or incompletely folded proteins from the ER in response to ER stress. Proteins targeted by ERAD are normally degraded via the ubiquitin-proteosome system (UPS) but in some circumstances may be degraded by the autophagy-lysosome system^58–60^. The two systems are also likely to be compensatory^59,61^. Consistent with this notion, SEL1L, a key component of mammalian ERAD^62^, was found to interact with MR1 in our system and MR1 degradation at steady state does not appear to be proteasome dependent, albeit at early timepoints^25,27^. However, further work would be required to explore this link in more detail.

Interestingly, both increased and decreased SQSTM expression (via depletion of Atg proteins and SQSTM1 respectively) results in increased activation of MAIT cells in the presence of *E.coli*. Thus, dysregulation of the autophagy pathway in general may be more important than the effects of individual proteins. Future work should aim to delineate the contributions of specific autophagy related proteins and the exact mechanisms at play.

Autophagy is crucial for cellular homeostasis and as demonstrated here and elsewhere, is an important regulator of antigen presentation, carefully controlling immune responses to pathogen challenge. This is evident by the fact that dysregulation of autophagy leads to numerous disease states including cancer, neurodegeneration and autoimmunity. Further work will be required to better understand the nature of autophagy mediated regulation of MR1 and the implications for both microbial control and recognition of cancerous cells.

## Materials and Methods

### Cell Culture

293T and HeLa cells were cultured in DMEM and Thp1 and C1R cells were cultured in RPMI both supplemented with 10% fetal bovine serum and penicillin-streptomycin. Thp1 and HeLa cells stably expressing a carboxyl-terminal hemaglutinin (HA) tagged MR1 were generated by standard lentivirus mediated transduction, followed by puromycin selection. Thp1 and/or C1R cells stably expressing either control, Atg5, Atg7, β2M or SQSTM1 targeting guide RNAs were generated by standard lentivirus mediated transduction followed by selection with puromycin or GFP. Clonal cell populations were generated by limiting dilution. Healthy donor blood leukocyte cones were obtained from NHS Blood and Transplant Service at the John Radcliffe Hospital, Oxford under GI Biobank ethics 16/YH/0247.

### CRISPR-Cas9

Guide RNAs were cloned into plentiCRISPRv2 (Addgene #52961) or plentiCRISPRv2GFP (Addgene #82416) using the BsmB1 restriction site. Cloned inserts were verified to be correct by sequencing. The sequences of the guide RNAs used in this study can be found in Supplementary Table S2.

### Fixed Bacteria

*E. coli* (DH5 alpha) were cultured overnight at 37°C in Luria broth (Sigma), washed in PBS and then fixed in 4% paraformaldehyde for 20 mins at room temperature. Bacteria were washed twice in PBS and resuspended in sterile filtered PBS. Bacteria were quantified as previously described^17^ and added at a concentration of 30 bacteria per cell (bpc) unless otherwise stated.

### Ligands

5-OP-RU was synthesized in DMSO as previously reported^6,7^, and used as a DMSO solution due to its much greater stability than in water, with care taken to minimize exposure time to water in cellular experiments. Ac-6-FP (Cayman Chemicals, 23303) was resuspended in DMSO and both synthetic ligands were used at a concentration of 10μM unless otherwise stated.

### Autophagy Inhibitors

5mM 3MA (Sigma, M9281) and 1μM Wortmannin (Sigma, 12338) were added to Thp1 or HeLa cells for 4 hours. For autophagy induction experiments, cells were washed in PBS and resuspended in EBSS media (Gibco) for 4 hours, with 100nM Bafilomycin (Sigma, 19-148) added for the final 2 hours.

### Coimmunoprecipitation

5x10^6^ Thp1 or Thp1.MR1.HA cells were incubated overnight (or 2 hours) with and without fixed *E. coli* or 5-OP-RU. Cells were harvested, washed in phosphate buffered saline (PBS) and resuspended in 500μL Lysis Buffer [0.5% NP40, 150 mM NaCl, 50 mM Tris-HCL (pH 7.5), protease inhibitor (+ EDTA)] for 30 mins at 4°C with rotation. Lysates were cleared by centrifugation for 10 mins at 1000xg with 20ul removed as the Input sample and the rest added to HA conjugated Dynabeads (ThermoFisher) for 2 hours at 4°C with rotation. The beads were washed 3 times, resuspended in loading buffer and protein eluted by boiling prior to immunoblot analysis.

### Immunoblots

For protein expression from cell lysates, cells were washed in PBS and resuspended in RIPA Buffer [50mM Tris-HCL (pH 8.0), 150mM NaCl, 1% NP40, 0.5% sodium deoxycholate, protease inhibitor (+ EDTA)] and incubated at 4°C for 20 mins with rotation. Lysates were cleared by centrifugation for 20 mins at 12,000xg and the supernatants transferred to a fresh tube. Total protein concentration was determined using the Pierce BCA Protein Assay kit (Fisher Scientific, 18064090), with 30-50ug of total protein loaded into each well of a 4-12% Tris-Glycine SDS-PAGE gradient gel (ThermoFisher Scientific). Resolved proteins were analyzed by immunoblotting using the following primary antibodies; anti-Atg5 (ab108327, Abcam), anti-Atg7 (8558, Cell Signalling), anti-AUP1 (35055, Cell Signalling), anti-Bag2 (ab79406, Abcam), anti-βactin (ab227387, Abcam), anti-β2M (ab75853, Abcam), anti-Calnexin (ab22595, Abcam), anti-Calreticulin (ab92516, Abcam), anti-GAPDH (ab8245, Abcam), anti-HA (NB600-363, Novus Biologicals), anti-Hsp90 (sc-13119, Santa Cruz Biotechnology), anti-IL23 (ab45420, Abcam), anti-LC3B (ab51520, Abcam), anti-SEL1L (ab78298, Abcam), anti-SQSTM1 (ab109012, Abcam), anti-STING (ab181125, Abcam), anti-Tapasin (MABF249, Merck). Bound antibodies were visualized using appropriate horseradish peroxidase conjugated secondary antibodies and enhanced chemiluminscence. For EndoH experiments, following the final wash proteins were eluted in Glycoprotein Denaturing Buffer (NEB, P0702S) for 10 mins at 100°C and then resuspended in EndoH (NEB, P0702S) for 1.5 hours at 37°C. Samples were then prepared for immunoblotting as described.

### Immunofluorescence

Sub-confluent HeLa cell monolayers were seeded onto glass coverslips. Twenty-four hours later, cells were treated with the appropriate autophagy reagent, as described above, and then washed three times in PBS and fixed with 4% paraformaldehyde (Electron Microscopy Sciences) for 15 mins. Cells were permeabilized with 0.2% Triton X-100 and then blocked with NGB Buffer (50 mM NH4Cl, 1% goat serum, 1% bovine serum albumin) for 1 hour. Cells were then incubated with the indicated primary antibodies for 2 hours and the appropriate species specific Alexa Fluor 488 or 594 conjugated secondary antibody for 1 hour. Coverslips were mounted onto slides using Diamond Anti-Fade Mounting Reagent Plus DAPI (ThermoFisher). Images are maximum intensity z-stack projections acquired on an Olympus FV1000 laser scanning confocal microscope. Fields of view were determined based on DAPI expression, with at least three fields of view imaged per sample and the number of cells analysed noted in the figure legend. Quantitative analysis was performed using ImageJ software. MF1 was quantified for individual cells after segmentation using CellPose and background subtraction. A size and intensity threshold was applied to enumerate the number of SQSTM1 foci per cell.

### Proteomics/Mass Spec

10^8^ Thp1 and Thp1-MR1 cells per condition were incubated with fixed *E.coli* overnight. Cells were washed and resuspended in Lysis Buffer (as above) and incubated for 30 mins at 4°C. Lysates were cleared by centrifugation for 10 mins at 1000xg and added to HA conjugated Dynabeads for 2 hours with rotation. The beads were washed 4 times in Wash Buffer (with the final wash without detergent) and proteins were eluted from the beads using 2% acetic acid. Eluted samples were trypsin digested and run on a liquid chromatography tandem MS (LC-MS/MS) system. Peptides were resuspended in 5% formic acid and 5% DMSO and then trapped on a C18 PepMap100 pre-column (300µm i.d. x 5mm, 100 Å, Thermo Fisher Scientific) using solvent A (0.1% Formic Acid in water) at a pressure of 500 bar and separated on an Ultimate 3000 UHPLC system (Thermo Fischer Scientific) coupled to a QExactive mass spectrometer (Thermo Fischer Scientific). The peptides were separated on an in-house packed analytical column (360µm x 75µm i.d. packed with ReproSil-Pur 120 C18-AQ, 1.9µm, 120 Å, Dr.Maisch GmbH) and then electrosprayed directly into an QExactive mass spectrometer (Thermo Fischer Scientific) through an EASY-Spray nano-electrospray ion source (Thermo Fischer Scientific) using a linear gradient (length: 60 minutes, 15% to 35% solvent B (0.1% formic acid in acetonitrile), flow rate: 200 nL/min). The raw data was acquired on the mass spectrometer in a data-dependent mode (DDA). Full scan MS spectra were acquired in the Orbitrap (scan range 350-2000 m/z, resolution 70000, AGC target 3e6, maximum injection time 50 ms). After the MS scans, the 20 most intense peaks were selected for HCD fragmentation at 30% of normalized collision energy. HCD spectra were also acquired in the Orbitrap (resolution 17500, AGC target 5e4, maximum injection time 120 ms) with first fixed mass at 180 m/z. The raw data files generated were processed using MaxQuant (Version 1.5.0.35), integrated with the Andromeda search engine. For protein groups identification, peak lists were searched against human database (Swiss--Prot, version 04/13) as well as a list of common contaminants by Andromeda. Trypsin with a maximum number of missed cleavages of 2 was chosen. Acteylation (Protein N—term) and Oxidation (M) were selected as variable modifications while Carbamidomethylation (C) was set as a fixed modification. Protein and PSM false discovery rate (FDR) were set at 0.01.

### Flow Cytometry

For the MAIT cell activation assay, Thp1 cells were incubated with or without fixed *E. coli* or 5-OP-RU overnight. PBMCs from healthy donors (NHS Blood Bank, Oxford) were thawed and CD8+ T cells positively enriched using CD8 microbeads (Miltenyi Biotech) as per the manufacturer’s instructions. CD8+ T cells were incubated with Thp1 cells at a 1:1 ratio for 5 hours, with Brefeldin A added for the final 4 hours. Cells were washed with PBS and stained with Live/Dead Fixable Near IR dye (Invitrogen) for 20 mins at room temperature. Cells were washed and fixed with 2% paraformaldehyde for 20 mins at 4°C. Cells were then washed twice in 1X Permeabilization Buffer (eBioscience) and stained with the following antibodies; CD3-Pe-Cy7 (25-0038-42, eBioScience), CD4-Viogreen (130-113-230, Miltenyi Biotech), CD8-eFluor450 (8-0086-42, Invitrogen), CD161-PE (130-113-593, Miltenyi Biotech), Vα7.2-APC (351708, Biolegend) and IFNγ-PerCpCy5.5 (45-7319-42, eBioScience). Cells were incubated for 25 mins at room temperature, washed and acquired on a MACSQuant Cytometer (Miltenyi Biotech).

For MR1 surface staining, cells were incubated with or without fixed *E. coli*, 5-OP-RU or Ac-6-FP overnight or for up to 6 hours. For cyclohexamide (CHX) experiments, cells were first incubated with 10μM CHX (Merck) for 1 hour before the addition of ligand for 6 hours. Cells were washed with PBS and then incubated with Human TruStain FcX Blocking Solution (Biolegend) for 10 mins. Cells were stained with MR1-PE (361106, Biolegend) or a matched isotype control (400214, Biolegend) antibody for 20 mins at room temperature. Cells were fixed in 2% paraformaldehyde for 20 mins at 4°C, washed in PBS and acquired on a MACSQuant. All data were analyzed using FlowJo version 10 software. For MHC-I surface staining, cells were stained with an HLA-ABC-FITC (11-9983-42, eBioScience) antibody.

For monocyte staining, PBMCs from healthy donors were first CD3 depleted using CD3 microbeads (Milteny Biotech) as per the manufacturer’s instructions and then stained using CD14-FITC (301804, Biolegend), CD16-PeCy7 (980110, Biolegend) and HLADR-BV41 (307636, Biolegend).

### qRT-PCR

RNA was extracted from Thp1 cells using the RNeasy mini kit (Qiagen), as per the manufacturer’s instructions, and equal volumes reverse transcribed using the appSCRIPT cDNA synthesis kit (Appleton Woods). qPCRs were performed in triplicate using Universal ProbeLibrary probes (Roche) on a LightCycler 480 machine (Roche). Results were normalized to the housekeeping genes GAPDH and HRPT. The sequences of the primers used can be found in Supplementary Table S2.

## Statistical Analysis

Statistical significance was calculated by either a one or two way ANOVA or a two tailed paired t-test, as indicated in the figure legends, using GraphPad Prism software (version 7).

## Data availability

The proteomics dataset generated during the current study has been deposited to the ProteomeXchange Consortium via the MassIVE repository with the dataset identifier MSV000089910 and is available here: https://massive.ucsd.edu/ProteoSAFe/privatedataset.jsp?task=7f9ad709f11a4ba88ea03dc43050a17a.

## Acknowledgements

We wish to thank the Advanced Proteomics Facility, University of Oxford, and the Micron Bioimaging Facility, University of Oxford, for their assistance with this work. We would also like to thank Dr. Matthew Edmans for careful reading of the manuscript and Drs Lucy Garner and Narayan Ramamurthy for helpful discussions. D.P.F. is supported by funding from the National Health and Medical Research Council (2009551), and the National Institute of Allergy and Infectious Diseases of the National Institutes of Health (R01AI148407NIH). P.K. is supported by funding from the Wellcome Trust (109965MA and 222426/Z/21/Z).

## Author contributions

Conceptualization and Methodology, P.P., J.U. and P.K.; Investigation, P.P., S.H., and C-P.H.; Formal Analysis, P.P., S.H., E.M.; Resources, J.U., J.Y.M.K. and D.P.F.; Writing-Original Draft, P.P.; Writing-Review and Editing, P.P., J.U., J.Y.M.K., D.P.F. and P.K.; Supervision and Funding Acquisition, P.K.

