## Supplementary Data for "MR1 dependent MAIT cell activation is regulated by autophagy associated proteins"

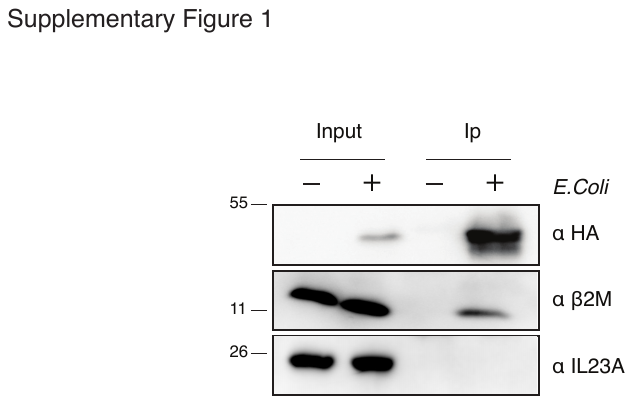

**Supplementary Figure S1:** Thp1.MR1.HA cells incubated with or without fixed *E.coli* overnight were lysed and MR1 immunoprecipitated using a HA antibody conjugated to magnetic beads. Immunoprecipitates (Ip) and Input samples were analyzed by western blot and probed with the indicated antibodies. The position of molecular weight markers bands (kDa) are indicated on the left. Immunoblot is representative of 3 independent experiments.

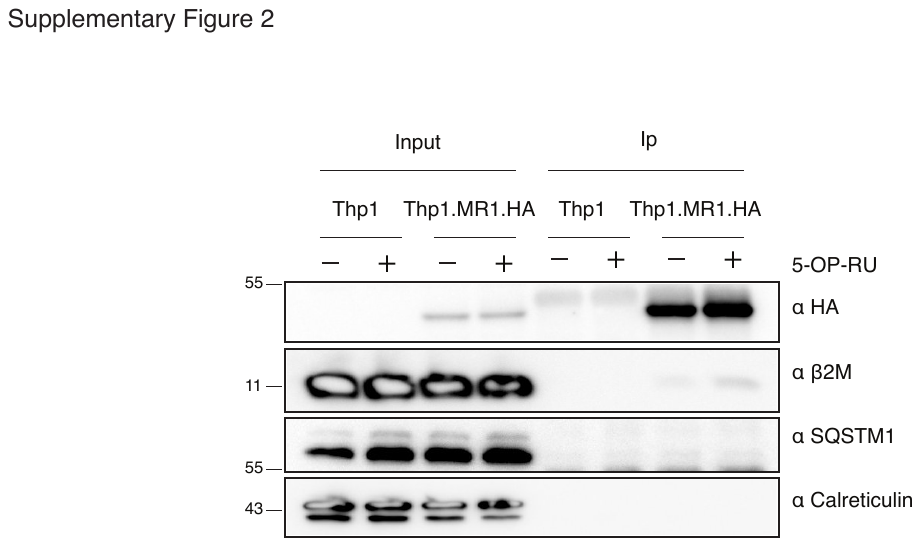

**Supplementary Figure S2:** Thp1 and Thp1.MR1.HA cells incubated with 5-OP-RU for 2 hours were lysed and MR1 immunoprecipitated using a HA antibody conjugated to magnetic beads. Ip and input samples were analyzed by western blot and probed with the indicated antibodies. The position of molecular weight markers bands (kDa) are indicated on the left. Immunoblot is representative of 3 independent experiments.

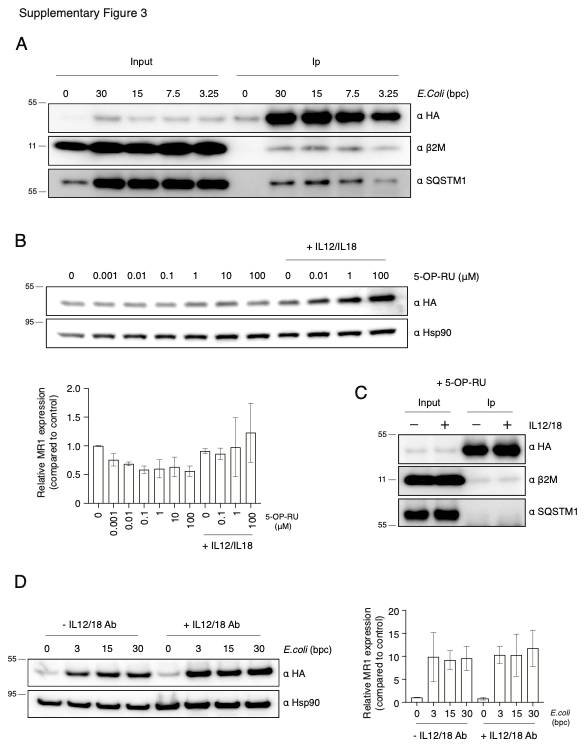

**Supplementary Figure S3:** A**.** Thp1.MR1.HA cells incubated with a titration of *E.coli* overnight were lysed and MR1 immunoprecipitated with a HA antibody conjugated to magnetic beads. Ip and Input samples were analyzed by western blot and probed with the indicated antibodies. bpc = bacteria per cell. B. Thp1.MR1.HA cells incubated with a titration of 5-OP-RU in the presence and absence of 50ng/mL IL12 and IL18 cytokines. Cells were lysed and analyzed by western blot by probing with the indicated antibodies. Quantification of the western blot is displayed in the bar chart. Data are the average from 2 independent experiments. C. Thp1.MR1.HA cells incubated with 5-OP-RU overnight, in the presence and absence of 50ng/mL IL12 and IL18 cytokines, were lysed and MR1 immunoprecipitated with a HA antibody conjugated to magnetic beads. Ip and Input samples were analyzed by western blot and probed with the indicated antibodies. D. Thp1.MR1.HA cells were incubated with *E.coli* and IL12 and IL18 blocking antibodies or isotype controls overnight. Cells were lysed and analyzed by western blot by probing with the indicated antibodies. The position of molecular weight markers bands (kDa) are indicated to the left of all immunoblots. Quantification of the western blot is displayed in the bar chart. Data are the average from 2 independent experiments All data are representative of at least 2 independent experiments.

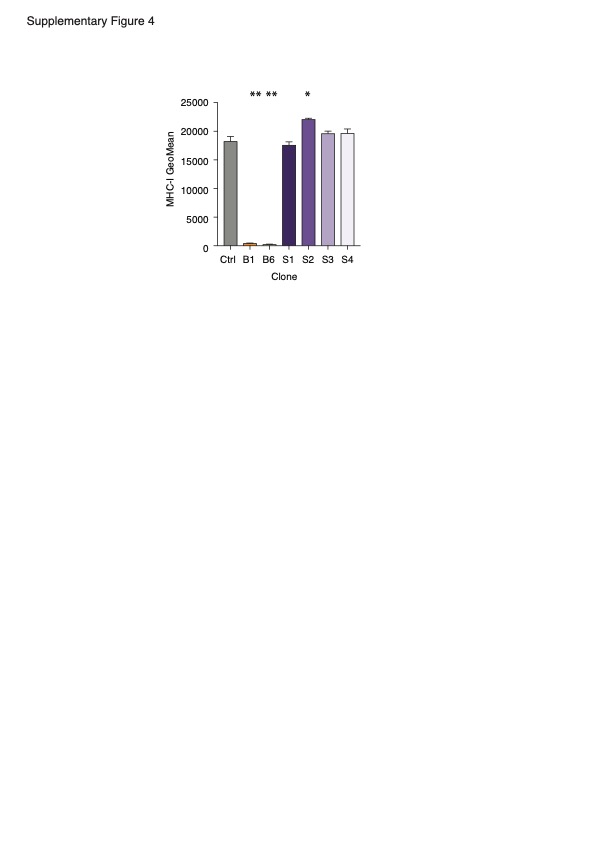

**Supplementary Figure S4**: Bar graph showing geometric MFIs of MHC-I surface expression in B2M (B1 and B6) and SQSTM1 (S1-S4) CRISPR-Cas9 treated clonal populations of Thp1 cells. Displayed is the average ± SD of 2 independent experiments. Statistical significance was calculated using a one-way ANOVA, ** = p < 0.0005, * = p < 0.05.

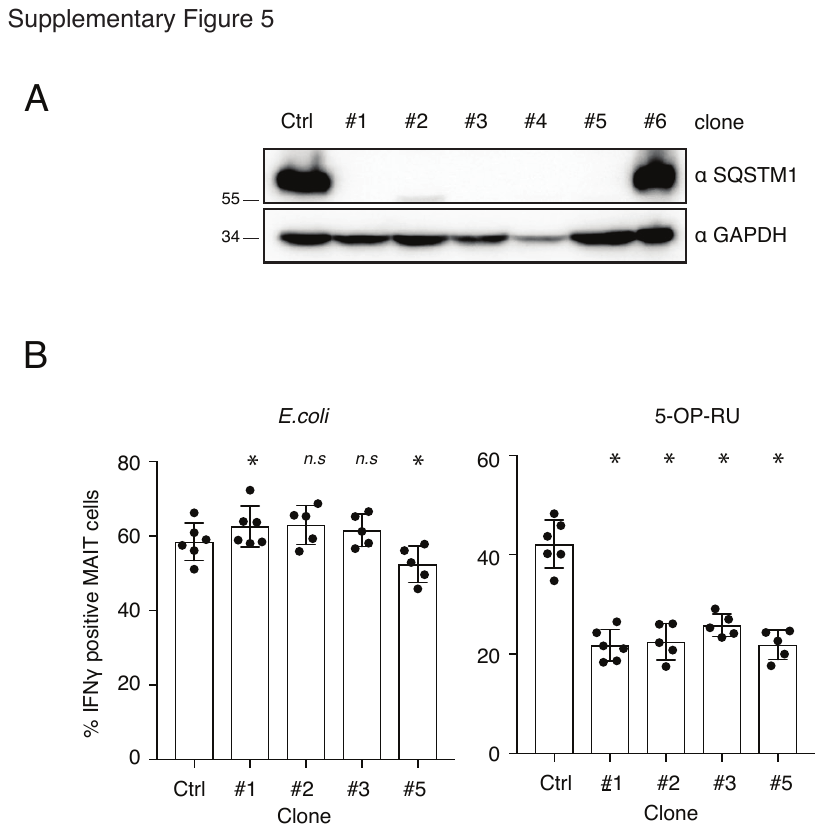

**Supplementary Figure S5**: A. Clonal populations of Thp1 cells treated with a guide RNA targeting SQSTM1 (clones #1-6) were established by limiting dilution. Cells were lysed and protein expression confirmed by western blot analysis by probing with the indicated antibodies. The position of molecular weight marker bands (kDa) are indicated on the left. B. Thp1 cells as described in A and control cells were incubated with fixed *E.coli* or 5-OP-RU overnight and then co-cultured with enriched CD8+ cells for 5 hours. IFNγ production ± SD from the CD161++Vα7.2+ population was measured by FACS. Data were acquired from 6 donors across 2 experiments although for some samples one of the donors had to be excluded due to a technical error. Statistical significance was calculated using a one-way ANOVA, * = p < 0.05, ns = p > 0.05.

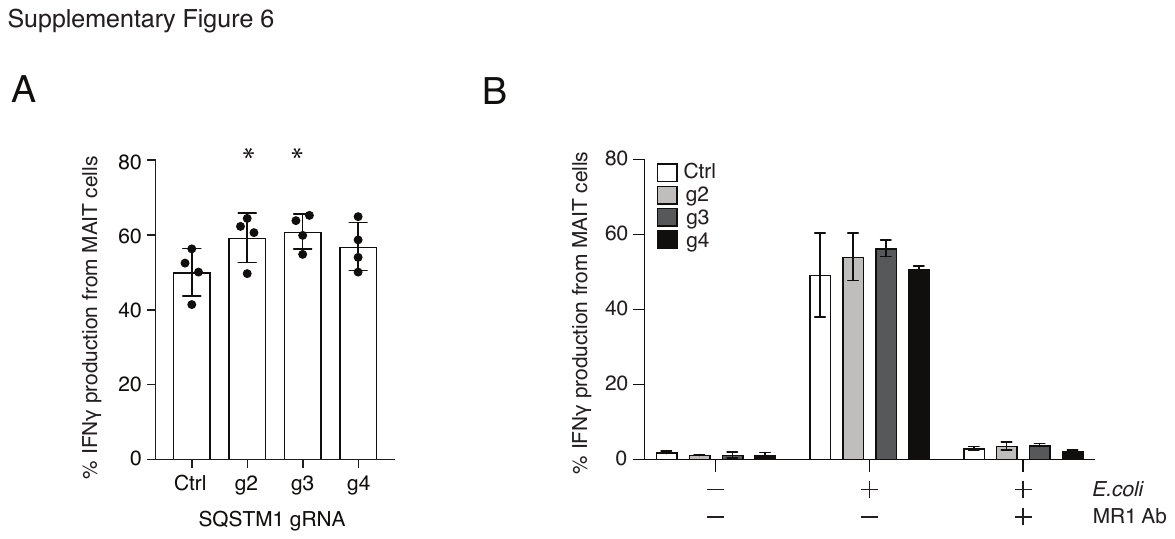

**Supplementary Figure S6**: A. Thp1 cells transduced with guide RNAs targeting SQSTM1 (g2, g3 and g4) or control cells (Ctrl) were incubated with fixed *E.coli* overnight and then co-cultured with enriched CD8+ cells for 5 hours. IFNγ production ± SD from the CD161++Vα7.2+ population was measured by FACS. Data were acquired from 4 donors across 2 experiments. B. SQSTM1 depleted Thp1 cells, as described in A, were incubated with or without *E.Coli* overnight and then co-cultured with enriched CD8+ cells for 5 hours in the presence and absence of an MR1 blocking antibody. IFNγ production ± SD from the CD161++Vα7.2+ population was measured by FACS. Data were acquired from 2 donors across 2 experiments.

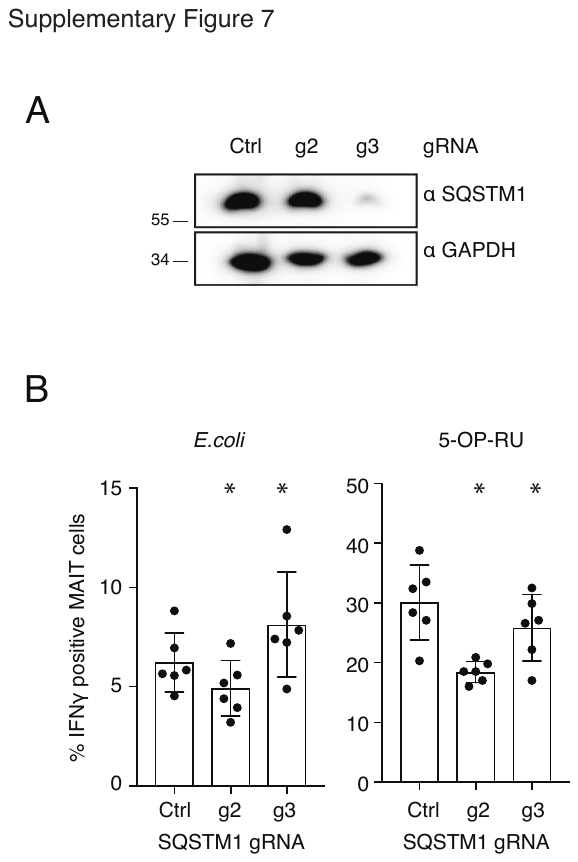

**Supplementary Figure 7**: A. C1R cells transduced with guide RNAs targeting SQSTM1 or control cells were lysed and analyzed by western blot by probing with the indicated antibodies. The position of molecular weight marker bands (kDa) are indicated on the left. B. C1R cells as described in A were incubated with fixed *E.coli* or 5-OP-RU overnight and then co-cultured with enriched CD8+ cells for 5 hours. IFNγ production ± SD from the CD161++Vα7.2+ population was measured by FACS. Data were acquired from 6 donors across 2 experiments. Statistical significance was calculated using a one-way ANOVA, * = p < 0.05, ns = p > 0.05.

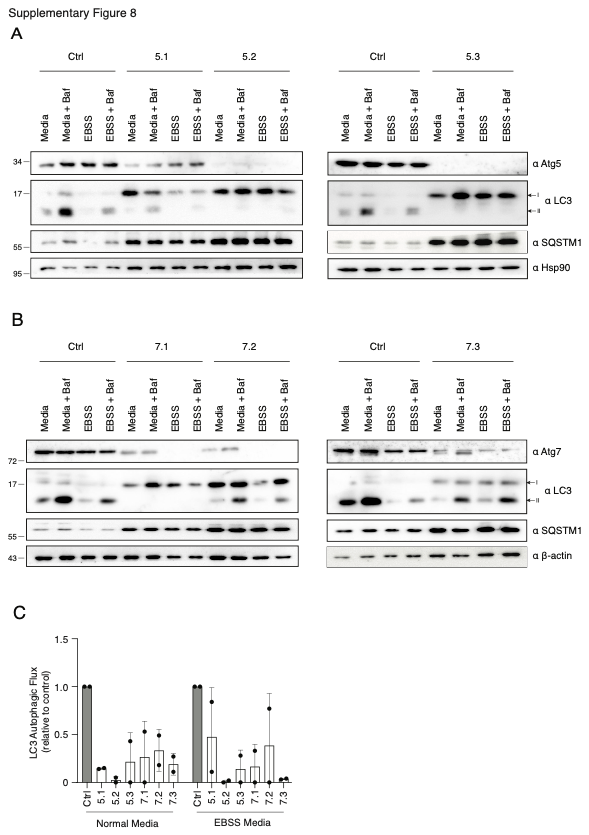

**Supplementary Figure S8:** Thp1 cells transduced with guide RNAs targeting Atg5 (A) or Atg7 (B) or control cells were incubated with normal media or EBSS media for 2 hours before addition of bafilomycin for an additional 2 hours. Cells were lysed and analyzed by western blot by probing with the indicated antibodies. The LC3B-I (I) and LC3B-II (II) bands are indicated to the right and the position of molecular weight marker bands (kDa) are indicated on the left of all immunoblots. Quantification of the western blots and determination of LC3 autophagic flux is displayed in the bar chart. Data is the average of 2 independent experiments.

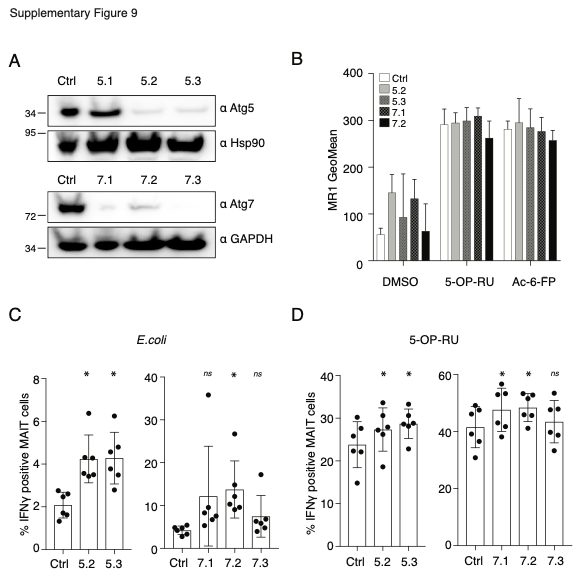

**Supplementary Figure S9:** A. C1R cells transduced with guide RNAs targeting Atg5 (top) and Atg7 (bottom) and control cells were lysed and knock down efficiency determined by western blot by probing with the indicated antibodies. The position of the molecular weight marker bands (kDa) are indicated on the left. B. Bar plot showing MR1 surface expression from the C1R cells described in A following overnight incubation with DMSO, 5-OP-RU or Ac-6-FP. Displayed is the average geometric MFI ± SD from 3 independent experiments. C1R cells as described in A were incubated overnight with fixed *E.coli* (C) or 5-OP-RU (D) and then co-cultured with enriched CD8+ cells for 5 hours. IFNγ production from the CD161++Vα7.2+ population was measured by FACS. For C and D data were acquired from 6 donors across 2 experiments. Statistical significance was calculated using a one-way ANOVA, * = p < 0.05, ns = p > 0.05.

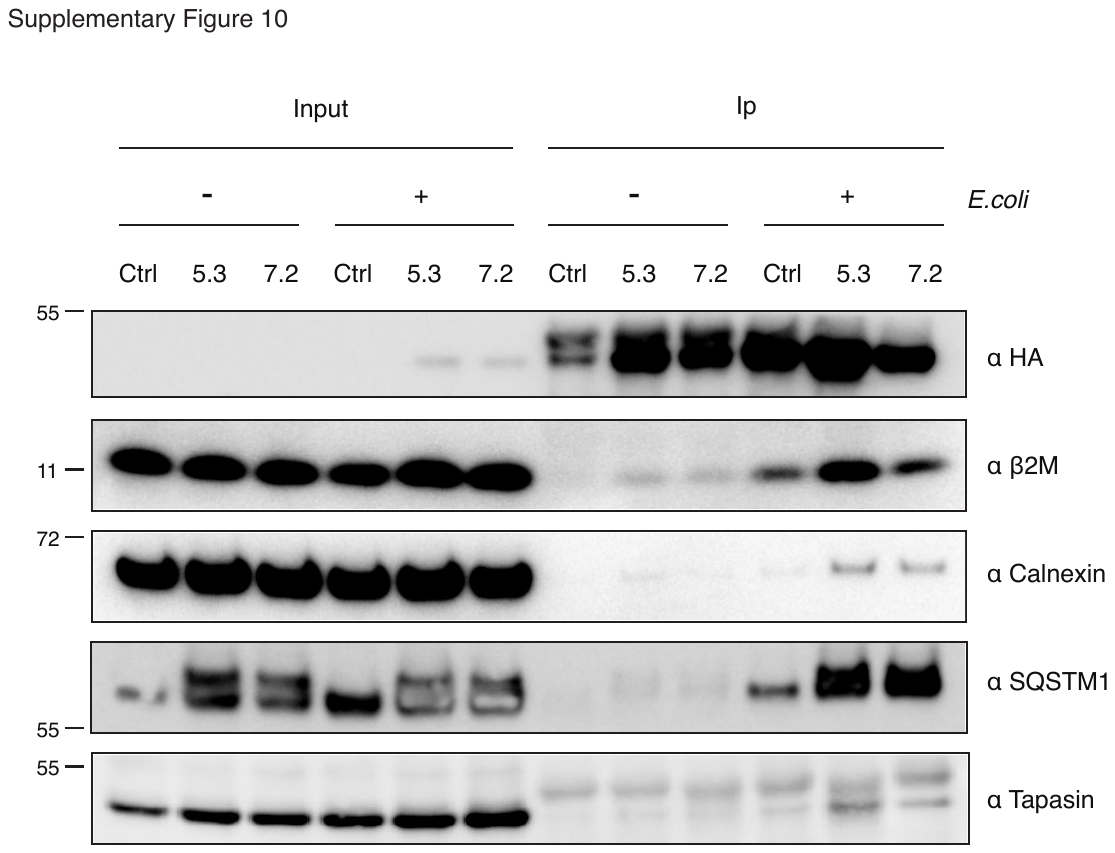

**Supplementary Figure S10:** Thp1.MR1.HA cells transduced with guide RNAs targeting Atg5 (5.3) and Atg7 (7.2) and control cells were incubated with fixed *E.coli* overnight and then lysed in 0.5% NP40. MR1 was immunoprecipitated using a HA antibody conjugated to magnetic beads. Immunoprecipitates were analyzed by western blot and probed with the indicated antibodies. The position of molecular weight marker bands (kDa) are indicated on the left. Shown are representative blots from one of 2 independent experiments.

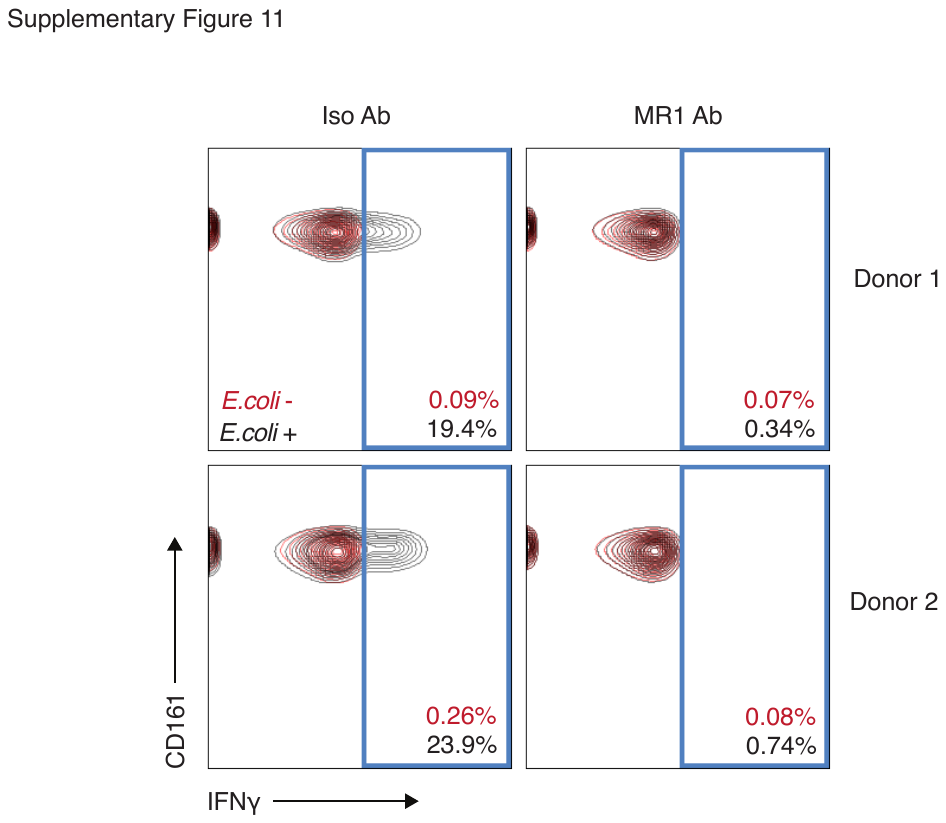

**Supplementary Figure S11:** HeLa.MR1.HA cells incubated with or without fixed *E.coli* overnight and co-cultured with enriched CD8+ cells from 2 donors for 5 hours in the presence or absence of an MR1 blocking antibody. IFNγ production from the CD161++Vα7.2+ population was measured by FACS and displayed in the contour plots.

**Supplementary Table S1**: Results from the proteomics screen showing proteins significantly enriched with MR1.HA (log fold change > 4) compared to the control.

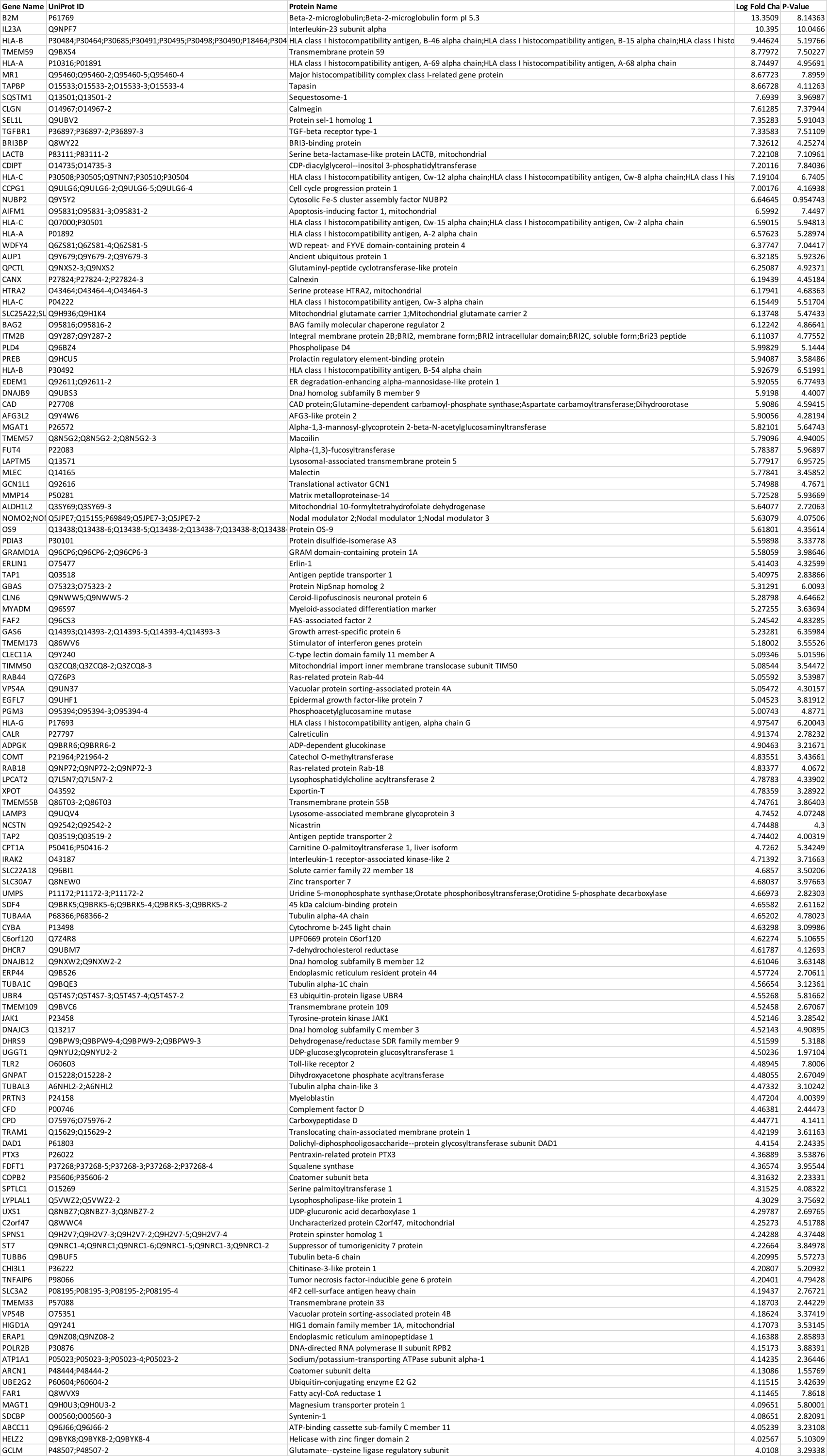

**Supplementary Table S2:** Oligonucleotides used in this study

|  | Forward | Reverse |
| --- | --- | --- |
| **CRISPR-Cas9 Cloning** | | |
| Atg5 guide 1 | caccgaacttgtttcacgctatatc | aaacgatatagcgtgaaacaagttc |
| Atg5 guide 2 | caacgaagagtaagttatttgacgt | aaacacgtcaaataacttactcttc |
| Atg5 guide 3 | caacgtgcttcgagatgtgtggtt | aaacaaccacacatctcgaagcac |
| Atg7 guide 1 | caccgaagctgaacgagtatcggc | aaacgccgatactcgttcagcttc |
| Atg7 guide 2 | caccgaactccaatgttaagcgagc | aaacgctcgcttaacattggagttc |
| Atg7 guide 3 | caccgaataatggcggcagctacgg | aaacccgtagctgccgccattatt c |
| β2M guide 5 | caccgactcacgctggatagcctcc | aaacggaggctatccagcgtgagtc |
| β2M guide 6 | caccgagtagcgcgagcacagcta | aaactagctgtgctcgcgctactc |
| SQSTM1 guide 2 | caccgttggggtgcaccatgttgcg | aaaccgcaacatggtgcaccccaac |
| SQSTM1 guide 3 | caccgcgagggaaagggcttgcac | aaacgtgcaagccctttccctcgc |
| SQSTM1 guide 4 | caccgaagatgtcatccttcacgt | aaacacgtgaaggatgacatcttc |
| Non-targeting control guide | caccgaagttcgagggcgacaccc | aaacgggtgtcgccctcgaacttc |
| **qPCR** | | |
| MR1 (Roche probe #19) | GCTGTCTCTGGGTCCATTGT | GATGGCTCCATTTTGCTCTC |
| GAPDH (Roche probe #60) | CCCCGGTTTCTATAAATTGAGC | CTTCCCCATGGTGTCTGAG |
| HRPT (Roche probe #73) | TGACCTTGATTTATTTTGCATACC | CGAGCAAGACGTTCAGTCCT |

Supplementary Methods

E.coli and 5OPRU Titration with IL12/18

Thp1.MR1.HA cells were incubated overnight with the indicated concentrations of 5-OP-RU or *E.coli* in the presence of 50ng/ml IL12 and IL18 blocking antibodies (or isotype control) or 50ng IL12 and IL18 cytokines (or DMSO). Cells were lysed and prepared for western blotting as previously described and blotted with the indicated antibodies.

Autophagy inhibition in Atg depleted cells

Atg5 and Atg7 depleted Thp1 cells were incubated in RPMI (media) or EBSS for 4 hours with Bafilomycin (Baf) added for the final 2 hours. Cells were then lysed and prepared for western blotting as previously described and blotted with the indicated antibodies.
